# Evolutionary recent dual obligatory symbiosis among adelgids indicates a transition between fungus and insect associated lifestyles

**DOI:** 10.1101/2020.10.16.342642

**Authors:** Gitta Szabó, Frederik Schulz, Alejandro Manzano-Marín, Elena Rebecca Toenshoff, Matthias Horn

## Abstract

Adelgids (Insecta: Hemiptera: Adelgidae) form a small group of insects but harbor a surprisingly diverse set of bacteriocyte-associated endosymbionts, which suggest multiple replacement and acquisition of symbionts over evolutionary time. Specific pairs of symbionts have been associated with adelgid lineages specialized on different secondary host conifers. Using a metagenomic approach, we investigated the symbiosis of the *Adelges laricis/tardus* species complex containing betaproteobacterial (‘ *Candidatus* Vallotia tarda’) and gammaproteobacterial (*‘Candidatus* Profftia tarda’) symbionts. Genomic characteristics and metabolic pathway reconstructions revealed that *Vallotia* and *Profftia* are evolutionary young endosymbionts, which complement each other’s role in essential amino acid production. Phylogenomic analyses and a high level of genomic synteny indicate an origin of the betaproteobacterial symbiont from endosymbionts of *Rhizopus* fungi. This evolutionary transition was accompanied with substantial loss of functions related to transcription regulation, secondary metabolite production, bacterial defense mechanisms, host infection and manipulation. The transition from fungus to insect endosymbionts extends our current framework about evolutionary trajectories of host-associated microbes.

## Introduction

Plant-sap feeding insects harbor bacterial endosymbionts, which are of great importance in their host ecology and serve as a model for studying microbe-host relationships and genome evolution of host restricted bacteria (1–3). Adelgids (Insecta: Hemiptera: Adelgidae) live on Pinaceae conifers and feed on phloem sap or parenchyma cells (4,5). The group has nearly seventy species and is sister to the families of phylloxerans (Phylloxeridae) and aphids (Aphididae) within the suborder Sternorrhyncha (6). Some adelgid species, such as the balsam woolly adelgid *(Adelges piceae)* and the hemlock woolly adelgid *(A. tsugae)* are well-known forest pests and represent severe threats to firs and hemlocks (7,8).

Adelgids have a complex multigenerational life cycle, which typically involves sexual generations and an alternation between spruce *(Picea)*, which is the primary host, and another, secondary conifer host *(Abies, Pinus, Larix, Pseudotsuga*, or *Tsuga*). However, other adelgids reproduce asexually in all generations on either of the host genera (5).

Similarly to other plant-sap feeding insects, adelgids harbor maternally inherited bacterial symbionts within specialized cells, so-called bacteriocytes, which form a large bacteriome in the abdomen (9–14). Although the function of these bacterial partners remains largely unexplored, they are expected to provide essential amino acids and B vitamins scarce in the plant sap diet, similarly to obligate endosymbionts of other plant-sap feeding insects (2,15). Besides these obligate nutritional endosymbionts, non-essential facultative symbionts might also occur within the bacteriome (or in other tissues), which can provide selective fitness benefits to insects such as protection against parasites and fungal pathogens, increased heat tolerance, or expansion of host plant range (16–18). Similarly to obligate mutualists, facultative symbionts are usually maternally inherited, but can also spread horizontally within and between insect species via mating (19), parasites (20), and food source such as plant tissues (21). Few examples of newly emerged bacteriocyte-associated symbionts of herbivorous insects pinpoint their source from plant-associated bacteria, such as *Erwinia* in *Cinara* aphids (22), gut bacteria, such as cultivable *Serratia symbiotica* strains colonizing the gut of *Aphis* aphids (23), and other free-living bacteria such as a *Sodalis* strain (HS) isolated from human wounds and being akin to primary endosymbionts of *Sitophilus* weevils (24).

Interestingly, two types of bacteriocyte-associated symbionts have been identified in all populations and life stages of most adelgid species (9–14). These symbionts belong to at least six different lineages within the Gammaproteobacteria or the Betaproteobacteria. *A. tsugae* populations also contain a universal *Pseudomonas* symbiont free in the hemocoel together with the bacteriocyte-associated symbionts (9,10,25). Remarkably, specific pairs of symbionts correspond to distinct lineages of adelgids specialized to one of the five secondary host tree genera (10,14). A gammaproteobacterial symbiont lineage involving *‘Candidatus* Annandia adelgestsugas’ and *‘Candidatus* Annandia pinicola’ (hereafter *Annandia)*, is present in both *A. tsugae* and *Pineus* species, and was likely already associated with ancestral adelgids before diversification into the two major adelgid lineages, *Adelges* and *Pineus*, over 87 million years ago (5,9,10,14). Nevertheless, this putatively ancient symbiont lineage is missing from other adelgids, and the high diversity of symbionts within this small group of insects suggests an evolutionary history involving multiple acquisitions and replacement of bacterial partners (10,13,14). This is in sharp contrast to the case in the aphid sister group, where most species have tightly co-evolved with a single obligate symbiont, *Buchnera aphidicola*, for over 180 million years (2). In the case of adelgids, it has been postulated that loss of the ancestral nutritional symbiont lineage and repeated replacements of bacterial partners might be due to fluctuating selective pressure on essential symbiotic functions during evolution as a consequence of repeated emergence of host-alternating lifestyles and feeding on nutrient-rich parenchyma versus nutrient-poor phloem on the primary and secondary host trees, respectively (10,25).

To date, whole genome sequences of adelgid endosymbionts are available for only one species: the hemlock woolly adelgid, *A. tsugae*. Metabolic potential and genomic characteristics of *Annandia* resemble those of long-term obligate intracellular symbionts. However, *Annandia* has lost many genes in essential amino acid synthesis. The accompanying, evolutionary more recent, *Pseudomonas* symbiont can complement these missing capabilities and thus has a co-obligatory status in maintaining the symbiosis (25). In addition to this obligate dual endosymbiotic system, analysis of a genome fragment of a gammaproteobacterial symbiont (*‘Candidatus* Steffania adelgidicola’) of the *Adelges nordmannianae/piceae* species complex revealed a metabolically versatile, putatively evolutionary young endosymbiont in this adelgid lineage (12). Further genomic data on the symbionts would help to infer the history of association of adelgids with distinct bacterial groups.

Here, we investigate the bacterial symbionts of the *A. laricis/tardus* species complex using a metagenomic approach and ask what is the function and putative origin of the dual symbiosis in this lineage of adelgids. *A. laricis* and *A. tardus* are morphologically and genetically hardly distinguishable species of adelgids (4,26). They contain betaproteobacterial and gammaproteobacterial symbionts, *‘Candidatus* Vallotia tarda’ and *‘Candidatus* Profftia tarda*’* (hereinafter *Vallotia* and *Profftia)*,respectively. Both symbionts are rod-shaped and are located intermingled inside the same bacteriocytes. *Profftia-related* symbionts have only been found in larch-associated, while *Vallotia* symbionts occur in both larch and Douglas-fir-associated lineages of adelgids. Although host-symbiont co-speciation could not be fully resolved with confidence yet, the dual obligatory status of *Profftia* and *Vallotia* in the symbiosis seems to be possible given their common occurrence across different populations and life stages of adelgids (10,13).

Our results demonstrate that both bacteriocyte-associated symbionts are evolutionary recent partners of adelgids complementing each other’s role in essential amino acid biosynthesis. Notably, phylogenomic analyses revealed a close relationship of *Vallotia* with endosymbionts of *Rhizopus* fungi. Detailed analysis of genomic synteny and gene content indicated an evolutionary transition from fungus to insect symbiosis accompanied by a substantial loss of functions in the insect symbiont especially in transcription regulation, secondary metabolite production, host infection and manipulation.

## Materials and Methods

### Sampling

Spruce (*Picea*) branches with galls of adelgids (4) were collected near Klausen-Leopoldsdorf, Austria (Figure S1). Galls were stored at −80°C in the lab for subsequent genomic DNA isolation.

### DNA isolation

Before DNA isolation, adelgids were collected from the galls using teasing needles. The insects were washed twice in buffer A + EDTA solution (35mM Tris-HCl, 250 mM sucrose, 25 mM KCl, 10 mM MgCl2, 250 mM EDTA; pH 7.5) and were subsequently homogenized in fresh solution with a plastic pestle. The suspension was then sequentially filtered through 53 and 30 μm pore size meshes and 5 μm membrane syringe filters. Samples were centrifuged at 7,000 rpm for 5 min at 4°C and supernatants were discarded. Pellets were re-suspended in buffer A and centrifuged again at 7,000 rpm for 5 min at 4°C. This washing step was repeated once and pellets were next re-suspended in 1xTE buffer (10 mM Tris-HCl, 1 mM EDTA; pH 7.5). High-molecular-weight DNA was isolated by an SDS-based DNA extraction method using 1% cetyltrimethylammonium bromide and 1.5% polyvinylpyrrolidone in the extraction buffer (27). DNA samples were stored at −20°C.

### Sequencing and genome assembly

A paired-end library was sequenced on a HighSeq 2000 Illumina sequencer. Sequencing reads were quality filtered and trimmed with PRINSEQ (28), and were assembled with SPAdes v3.1 (29). Using a subset of 30 million read pairs, a single contig representing the circular *Profftia* chromosome was obtained with 52-fold coverage, while the assembly of the *Vallotia* genome remained fragmented probably due to repetitive sequences. To improve this assembly, reads were mapped on the *Profftia* genome using the Burrows-Wheeler Alignment (BWA) tool and the BWA-MEM algorithm (30), and matching sequences were removed from further analysis. A novel assembly with the remaining reads resulted in 14 contigs longer than 1000 bp.

These contigs were further analyzed against a custom protein database containing single-copy markers found in 99% of prokaryote genomes using blastx (31) and phylogenetic information of the best hits was assessed in Megan v4.70.4 (32). Ribosomal RNAs were inferred by RNAmmer (33). Based on these, eight contigs belonging to the *Vallotia* genome were identified. Seven contigs represent the *Vallotia* chromosome with ~212-fold coverage. In addition, a single contig obtained with 169 fold coverage corresponds to a putative circular plasmid of this endosymbiont. The remaining contigs, all shorter than 5,500 bp, were judged to belong to unrelated taxa, based on differences in GC content, coverage and taxonomic affiliation of best blastx hits in the NCBI non-redundant protein database (nr).

### Genome analysis

The putative origin of replication was identified with GenSkew (http://genskew.csb.univie.ac.at/). We used the ConsPred genome annotation pipeline for gene prediction and annotation (34). Genome annotations were curated with the help of the UniPro UGENE software (35). We identified pseudogenes by using the intergenic and hypothetical protein regions as queries in blastx searches against nr and the UniProt Swiss-Prot database with an e-value < 1e-3 confidence threshold. Pseudogenes were identified as remains of genes, which were either truncated (having a length < 80% of the reference) or were interrupted by internal stop codons and/or frameshift mutations. Pseudogene coordinates were set according to the best blast hit in the UniProt Swiss-Prot database, if applicable.

The presence of mobile genetic elements was inferred with blastn and blastx searches against the ISfinder database (36). Metabolic pathways were explored with the help of the Ecocyc, Biocyc, and Metacyc databases (37) and the Pathway Tools software (38). Orthologous proteins shared by the relevant genomes or unique to either of the symbionts were identified by using OrthoMCL with a 1e-5 e-value threshold (39). Distribution of the predicted proteins among the main functional categories was explored by using eggNOG-mapper v2 (40,41) with the DIAMOND sequence comparison option and a 1e-3 e-value threshold (42). Genome alignments were performed by Mauve (43). The *Vallotia* contigs were reordered using the chromosome and plasmid sequences of the closely related fungus endosymbiont, *Mycetohabitans rhizoxinica*, as references (accession numbers: FR687359 and FR687360, respectively). Collinear genomic regions and genome rearrangements were visualized by Circos based on single-copy shared genes (44). A close-up of syntenic regions was created by using the Easyfig tool (45). The assembled genomes have been submitted to European Nucleotide Archive under accession number [submission in process, to be added].

### Phylogenetic analyses

A phylogenomic approach was used to infer the phylogenetic positions of the endosymbionts. Protein sequences of closely related species within the Burkholderiales and the Enterobacteriales were collected from the Assembly database of NCBI. Single copy marker genes were identified by Phyla-AMPHORA (46) using the Brandon Seah (2014) Phylogenomics-tools (online: https://github.com/kbseah/phylogenomics-tools). Individual sets of genes were aligned with Muscle 3.8.31 (47). Poorly aligned positions were removed with Gblocks 0.91b (48) using default parameters apart from the following settings: allowed gap positions with half, the minimum length of a block was 5. Alignments of 108 and 45 proteins were concatenated and used for the calculation of phylogenetic trees for *Vallotia* and *Profftia*, respectively.

For both endosymbionts, we generated maximum likelihood trees with IQ-TREE after selecting the best-fit substitution models with ModelFinder as available at http://iqtree.cibiv.univie.ac.at (49–51). SH-like approximate likelihood ratio test (SH-aLRT) and ultrafast bootstrap support values, both based on 1000 iterations, were calculated (52). The best-fit models were LG+F+R5 and LG+R5 for the *Vallotia* and *Profftia* tree, respectively. Besides, Bayesian phylogenetic analyses were performed by MrBayes 3.2.7a (53) with the LG+I+G model and 4 gamma categories on the CIPRES Science Gateway v.3.3. web interface (54). Two runs, each with 4 chains were performed until convergence diagnostics fell below 0.01. A 50% majority consensus tree was created with a relative burn-in of 25%.

## Results and discussion

### *Vallotia* and *Profftia* are evolutionary young symbionts of adelgids

#### Intermediate states of genome reduction

The complete sequence of the *Profftia* chromosome had a length of 1,225,795 bp and a G+C content of 31.9% (Table 1). It encoded for 645 proteins, one copy of each rRNA, 35 tRNAs, and 10 ncRNAs. It had tRNAs and amino acid charging potential for all 20 standard amino acids. However, protein-coding sequences made up only 52.4% of the genome, and 21 pseudogenes indicated an ongoing gene inactivation.

**Table 1.**
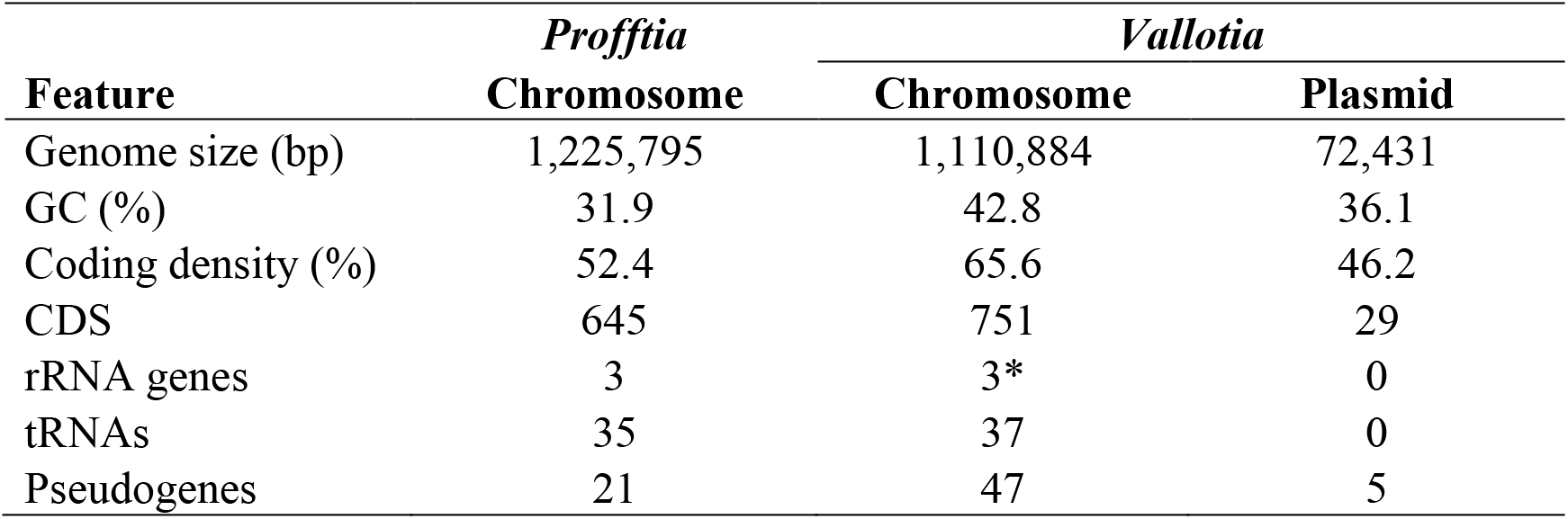
Genomic features of *Profftia* and *Vallotia*. *Vallotia likely has three copies of each rRNA gene based on the sequence coverage of the corresponding contigs involving these genes.

Currently, eight contigs with a total length of 1,183,315 bp represent the *Vallotia* genome. Seven contigs had an average G+C content and a coding density of 42.8% and 65.5%, respectively. However, a 72,431 bp long contig showed a characteristically lower G+C content (36.1%) and contained only 46.2% putative protein coding sequences (CDSs). This contig had identical repeats at its ends, and genome annotation revealed neighboring genes for a plasmid replication initiation protein, and ParA/ParB partitioning proteins, which function in plasmid and chromosome segregation between daughter cells before cell division (55). We thus assume that this contig corresponds to a circular plasmid of *Vallotia*. The 16S rRNA and the 5S plus 23S rRNA genes were encoded on two small contigs in the *Vallotia* assembly (1976 and 3571 bp, respectively) and were covered by nearly three times more sequence reads than the rest of the chromosomal contigs. This implies that *Vallotia* has three copies of each rRNA, similarly to its closest relative for which a complete genome sequence is available, *Mycetohabitans rhizoxinica* (56). In total, the *Vallotia* genome encoded for 780 proteins (29 on the putative plasmid), 37 tRNAs, and 52 predicted pseudogenes (5 on the putative plasmid).

The host-restricted lifestyle has a profound influence on bacterial genomes. Living in a stable, nutrient-rich niche relaxes purifying selection on many redundant functions, and small effective population size of the symbionts increases genetic drift. These can lead to the accumulation of slightly deleterious mutations, a proliferation of mobile genetic elements, and gene inactivation (57–60). Non-functional genomic regions and mobile genetic elements get subsequently lost, and ancient obligate endosymbionts typically have tiny (≪0.8 Mb), gene dense and stable genomes with AT-biased nucleotide composition (2,61,62). Facultative symbionts also possess accelerated rates of sequence evolution but have larger genomes (>2Mb) with variable coding densities following the age of their host-restricted lifestyle (63). The only moderately reduced size and AT bias together with the low protein-coding density of the *Vallotia* and *Profftia* genomes was most similar to those of evolutionary young co-obligate partners of insects (63), for instance ‘Ca. *Pseudomonas adelgestsugas’* in *A. tsugae* (25), *Serratia symbiotica* in *Cinara cedri* (64,65) and the *Sodalis-like* symbiont of *Philaenus spumarius*, the meadow spittlebug (66). However, compared to these systems involving a more ancient and a younger symbiont, similar genome characteristics of *Vallotia* and *Profftia* implied that both bacteria are evolutionarily recent symbionts in a phase of extensive gene inactivation typical for early stages of adaptation to an obligate host-restricted life-style (59,60).

#### Differential reduction of metabolic pathways

Although compared to their closest free-living relatives both *Vallotia* and *Profftia* have lost many genes in all functional categories, both retained a greater proportion of genes in translation-related functions and cell envelope biogenesis (Figures S2, S3). High retention of genes involved in central cellular functions such as translation, transcription, and replication is a typical feature of reduced genomes, even extremely tiny ones of long-term symbionts (62). However, ancient intracellular symbionts usually miss a substantial number of genes for the production of the cell envelope and might rely on host-derived membrane compounds (67–69).

Based on pathway reconstructions, both *Vallotia* and *Profftia* had a complete gene set for peptidoglycan, fatty acid, phospholipid biosynthesis, and retained most of the genes for the production of lipid A, LPS core, and the Lpt LPS transport machinery. Besides, we found a partial set of genes for O antigen biosynthesis in the *Vallotia* genome. Regarding the membrane protein transport and assembly, both adelgid endosymbionts had the necessary genes for Sec and signal recognition particle (SRP) translocation, and the BAM outer membrane protein assembly complex. *Profftia* also had a complete Lol lipoprotein trafficking machinery (lolABCDE), which can deliver newly matured lipoproteins from the inner membrane to the outer membrane (70). Besides, *Profftia* had a near-complete gene set for the Tol-Pal system, however, *tolA* has been pseudogenized suggesting an ongoing reduction of this complex. In addition, both adelgid endosymbionts have retained *mrdAB* and *mreBCD* having a role in the maintenance of cell wall integrity and morphology (71,72). The observed well-preserved cellular functions for cell envelope biogenesis and integrity are consistent with the rod-shaped cell morphology of *Profftia* and *Vallotia* (13), contrasting the spherical/pleomorphic cell shape of ancient endosymbionts, such as *Annandia* in *A. tsugae* and *Pineus* species (9,10,14).

Regarding the central metabolism, *Vallotia* lacks 6-phosphofructokinase, but has a complete gene set for gluconeogenesis and the tricarboxylic acid (TCA) cycle cycle. TCA cycle genes are typically lost in long-term symbionts but are present in facultative and evolutionary recent obligate endosymbionts (66,73,74). *Vallotia* does not have any sugar transporter genes, similarly to its close relative, the fungus symbiont *M. rhizoxinica* (56). A glycerol kinase gene next to a putative glycerol uptake facilitator protein is present on its plasmid, however, it has a frameshift mutation and a premature stop codon in the first 40% of the sequence and whether it can still produce a functional protein remains unknown.

*Profftia* can convert acetyl-CoA to acetate for energy but lacks TCA cycle genes, a feature characteristic to more reduced genomes, such, for instance, *Annandia* in *A. tsugae* (25). *Profftia* has import systems for a variety of organic compounds, such as murein tripeptides, phospholipids, thiamine, spermidine, and putrescine and 3- phenylpropionate and two complete phosphotransferase systems for the uptake of sugars. NADH dehydrogenase, ATP synthase, and cytochrome oxidases (*bo*/*bd*-1) are encoded on both adelgid symbiont genomes.

*Profftia* retained more functions in inorganic ion transport and metabolism, while *Vallotia* had a characteristically higher number of genes related to amino acid biosynthesis (see its function below) and nucleotide transport and metabolism (Figures S2, S3). For instance, *Profftia* can take up sulfate and use it for assimilatory sulfate reduction and cysteine production, and it has also retained many genes for heme biosynthesis. However it cannot produce inosine-5-phosphate (5′-IMP) and uridine 5′-monophosphate (5′-UMP) precursors for the *de novo* synthesis of purine and pyrimidine nucleotides thus would need to import these compounds.

Taken together their moderately reduced, gene-sparse genomes but still versatile metabolic capabilities support that *Vallotia* and *Profftia* are both in an intermediate stage of genome erosion and functional reduction similar to evolutionary recently acquired endosymbionts.

### *Vallotia* and *Profftia* are both obligatory nutritional symbionts

#### Complementary functions in essential amino acid provision

*Vallotia* and *Profftia* complement each other’s role in the essential amino acid synthesis, thus have a co-obligatory status in the *A. laricis/tardus* symbiosis (Figure 1). Although *Vallotia* likely generates most essential amino acids, it can not provide phenylalanine and tryptophan on its own. Solely, *Profftia* can produce chorismate, a key precursor for the synthesis of both amino acids. *Profftia* is likely responsible for the complete biosynthesis of phenylalanine as it has a full set of genes for this pathway. It can also convert chorismate to anthranilate, however further genes for tryptophan biosynthesis are only present in the *Vallotia* genome. Thus *Vallotia* likely takes up anthranilate for tryptophan biosynthesis. Anthranilate synthase (trpEG), is a subject to negative feedback regulation by tryptophan (75), thus partition of this rate-limiting step between the co-symbionts can enhance overproduction of the amino acid and might stabilize dual symbiotic partnerships at an early stage of coexistence. The production of tryptophan is partitioned between *Vallotia* and *Profftia* similarly as seen in other insect symbioses such as between *Buchnera* and *Serratia symbiotica* in co-obligatory partnerships in aphids (64,65) and between *Carsonella* and certain cosymbionts in psyllids (76). The tryptophan biosynthesis is also shared but is more redundant between the *Annandia* and *Pseudomonas* symbionts of *A. tsugae* (25). This association generally shows a higher level of functional overlap between the symbionts than the *Vallotia* - *Profftia* system, as redundant genes are present also in phenylalanine, threonine, lysine, and arginine synthesis in the *A. tsugae* symbiosis. Besides, the *Vallotia* - *Profftia* consortium is also more unbalanced than the *A. tsugae* system where *Annandia* can produce seven and the *Pseudomonas* partner five essential amino acids with the contribution of host genes (25).

**Figure 1.**
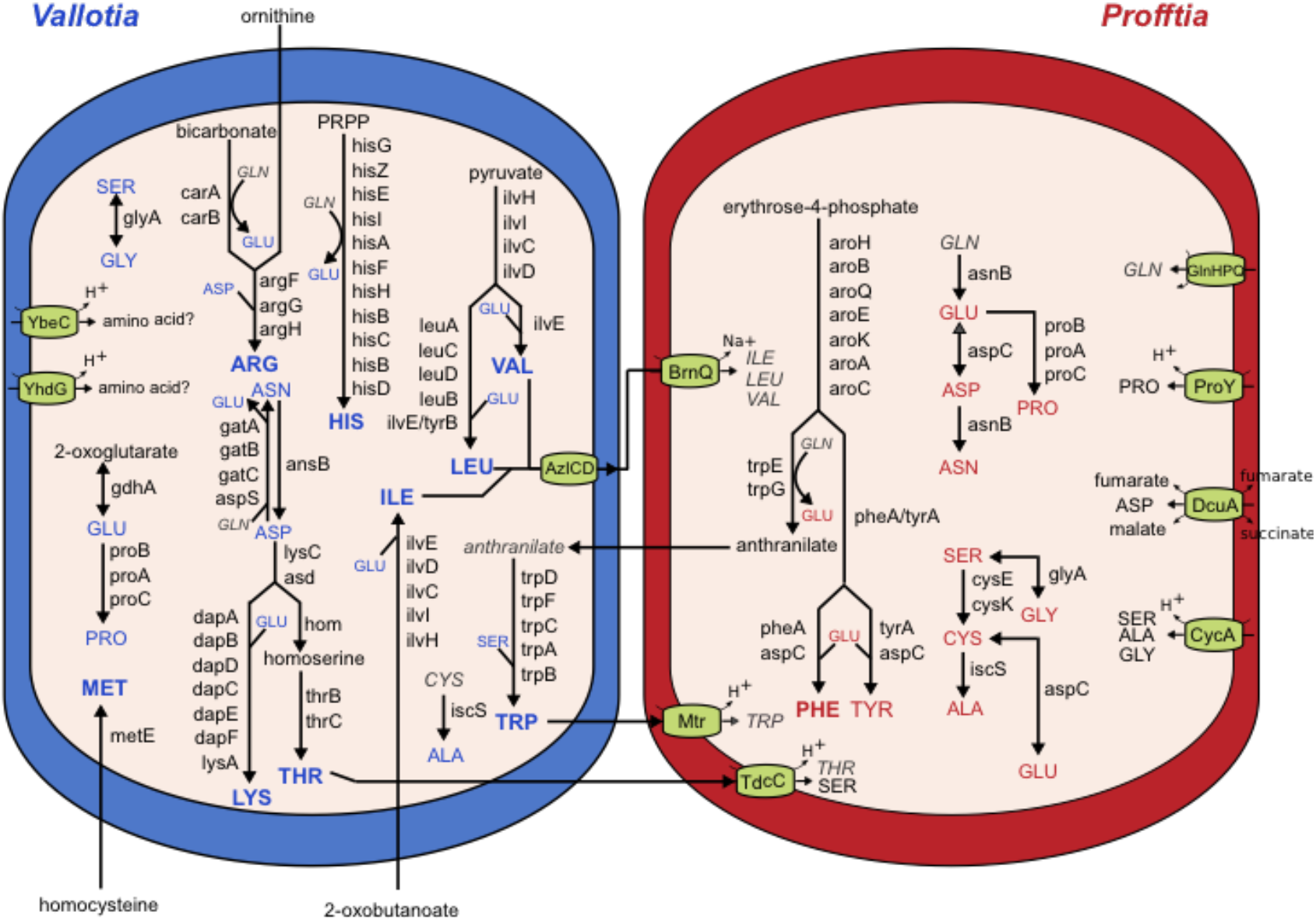
Division of labor in amino acid biosynthesis and transport between *Vallotia* and *Profftia* showing co-obligatory status of endosymbionts of *A. laricis/tardus.* Amino acids produced by *Vallotia* and *Profftia* are shown in blue and red, respectively. Bolded texts indicate essential amino acids. The insect host likely supplies ornithine, homocysteine, 2-oxobutanoate and glutamine. Other compounds that cannot be synthesized by the symbionts are shown in grey italics.

The *Vallotia* genome encodes for all the enzymes for the synthesis of five essential amino acids (histidine, leucine, valine, lysine, threonine). Among the essential amino acid synthesis related genes, *argG* and *tyrB* of *Vallotia* are only present on the plasmid, which might have contributed to its maintenance in the genome. However, neither of the endosymbionts can produce ornithine, 2- oxobutanoate and homocysteine *de novo*, which are key for the biosynthesis of arginine, isoleucine and methionine, respectively. *metC* and *argA* are still present as pseudogenes in *Vallotia* suggesting a recent loss of these functions in methionine and arginine biosynthesis, respectively. The corresponding functions are also missing from the *Annandia* - *Pseudomonas* system (Weglarz et al., 2018). Ornithine, 2- oxobutanoate, and homocysteine are thus likely supplied by the insect host, as seen for instance in aphids, mealybugs, and psyllids, where the respective genes encoding for cystathionine gamma and beta lyases and insect ornithine aminotransferase are present in the insect genomes and are typically overexpressed within the bacteriome (68,77,78). However, we can not confirm the presence of relevant genes in the *A. laricis/tardus* genome, as our metagenome data were almost free from eukaryotic sequences.

*Vallotia* and *Profftia* have more redundant functions in non-essential amino acid production. Both symbionts can synthesize seven non-essential amino acids mostly through a series of amino-acid conversions (Figure 1). Only *Profftia* can produce cysteine and tyrosine, while none of the symbionts can build up glutamine thus this latter amino acid is likely supplied by the insect bacteriocytes.

The presence of amino acid transporters can complement missing functions in amino acid synthesis in the endosymbionts (Figure 1). For instance *Profftia* has a high-affinity glutamine ABC transporter, and three symporters (BrnQ, Mtr, TdcC), which can import isoleucine, leucine, valine, tryptophan, and threonine among the essential amino acids that can be produced by *Vallotia*. *Vallotia* might excrete isoleucine, valine, and leucine via AzICD, a putative branched-chain amino acid efflux pump (79), and these amino acids could be taken up by *Profftia* via BrnQ and would be readily available also for the insect host.

#### B vitamin provision by Vallotia

Regarding the B vitamin synthesis, *Vallotia* should be able to produce thiamine (B1), riboflavin (B2), pantothenate (B5), pyridoxine (B6), biotin (B7), and folic acid (B9) (Figure S4). Although *Vallotia* misses some genes of the canonical pathways, alternative enzymes and host-derived compounds might bypass these reactions, as detailed in the supplementary material. *Profftia* has only a few genes related to B vitamin biosynthesis. Three pseudogenes *(ribAEC)* in the riboflavin synthesis pathway indicate that these functions might have been lost evolutionary recently in this symbiont.

In summary, *Profftia* and *Vallotia* are both obligate nutritional endosymbionts of adelgids, however, *Vallotia* has a pivotal role in essential amino acid and B vitamin provision.

### *Profftia* and *Vallotia* are related to free-living bacteria and fungus endosymbionts

Previous 16S rRNA-based phylogenetic analyses suggested an affiliation of *Profftia* with free-living gammaproteobacteria and a close phylogenetic relationship between *Vallotia* and betaproteobacterial endosymbionts of *Rhizopus* fungi (13). Biased nucleotide composition and accelerated sequence evolution of endosymbiont genomes (2,3) often result in inconsistent phylogenies and may cause grouping of unrelated taxa (68,80,81). Thus to further investigate the phylogenetic relationships of the *A. laricis/tardus* symbionts, we used conserved marker genes for maximum likelihood and Bayesian phylogenetic analyses.

Phylogenetic analysis of 45 single-copy proteins demonstrated that *Profftia* opens up a novel insect symbiont lineage most similar to *Hafnia* species and an isolate from the human gastrointestinal tract within the Hafniaceae, which has been recently designated as a distinct family within the Enterobacteriales (82) (Figure S5). *Hafnia* strains are frequently found in the gastrointestinal tract of humans and animals including insects, among others (83,84). The phylogenomic placement of *Profftia* in our analysis is in agreement with previous 16S rRNA based analyses (13).

*Vallotia* formed a monophyletic group with *Mycetohabitans endofungorum* and *M. rhizoxinica*, endosymbionts of *Rhizopus* fungi within the Burkholderiaceae (85,86) with strong support in phylogenetic analyses based on a concatenated set of 108 proteins (Figures 2, S6; previous taxonomic assignments of the fungus endosymbionts were as *Burkholderia/Paraburkholderia endofungorum* and *rhizoxinica*, respectively). Interestingly, *Vallotia* and *M. endofungorum* appeared as well-supported sister taxa within this clade. This implies a closer phylogenetic relationship between *Vallotia* and *M. endofungorum*, and a common origin of adelgid endosymbionts from within a clade of fungus endosymbionts. Lengths of branches leading to the fungus endosymbionts were similar to those of free-living bacteria in the data set, however *Vallotia* had a remarkably longer branch marking a rapid rate of sequence evolution characteristic of obligate intracellular bacteria (2,3). *M. endofungorum* and *M. rhizoxinica* have been identified in the cytosol of the zygomycete *Rhizopus microsporus*, best known as the causative agent of rice seedling blight (86,87). The necrotrophic fungus secrets potent toxins, rhizoxin and rhizonin, which are produced by the endosymbionts (86,88). The bacterial partners are obligatory for their host as they tightly control its sporulation, while they benefit from host nutrients and spread with the fungal spores (89,90). Additionally, related bacterial strains have also been found in association with *Rhizopus* fungi worldwide in a diverse set of environments, including other plant species, soil, food, and even human tissues (91–93).

**Figure 2.**
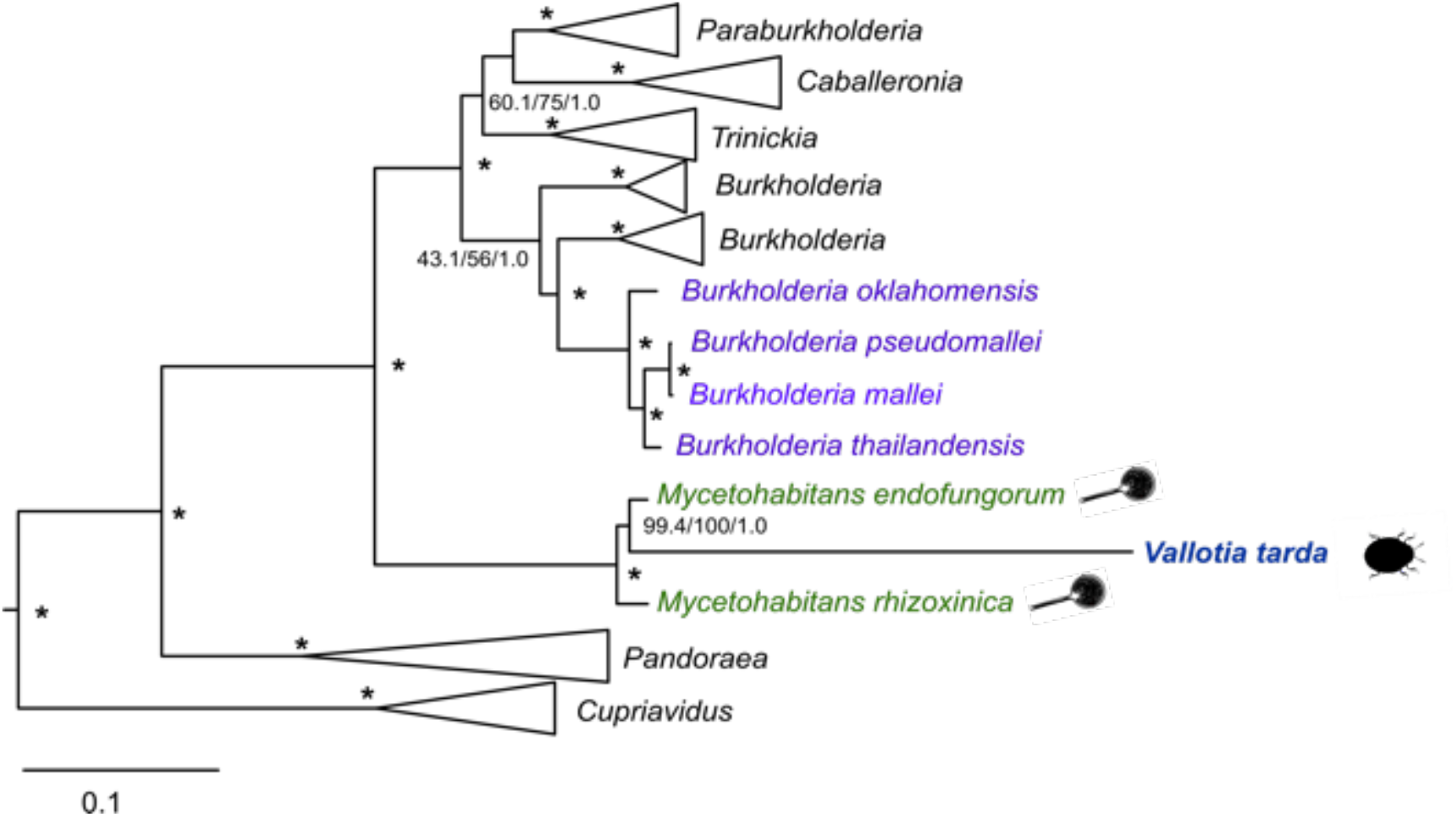
Phylogenomic analysis showing the affiliation of the adelgid endosymbiont *‘Candidatus* Vallotia tarda’ and its closest relatives, the fungus endosymbionts *M. rhizoxinica* and *M. endofungorum* within the *Burkholderiaceae.* Free-living and pathogenic bacteria are colored in purple. Selected members of *Oxalobacteraceae (Janthinobacterium agaricidamnosum* [HG322949], *Collimonaspratensis* [CP013234] and *Herbaspirillum seropedicae* [CP011930]) were used as outgroup. Maximum likelihood (IQ-TREE) and Bayesian analyses (MrBayes) were performed based on a concatenated alignment of 108 proteins. Maximum likelihood tree is shown. SH-aLRT support (%) and ultrafast bootstrap support (%) values based on 1000 replicates, and Bayesian posterior probabilities are indicated on the internal nodes. Asterisks stand for a maximal support in each analysis (100%/1).

**Figure 3.**
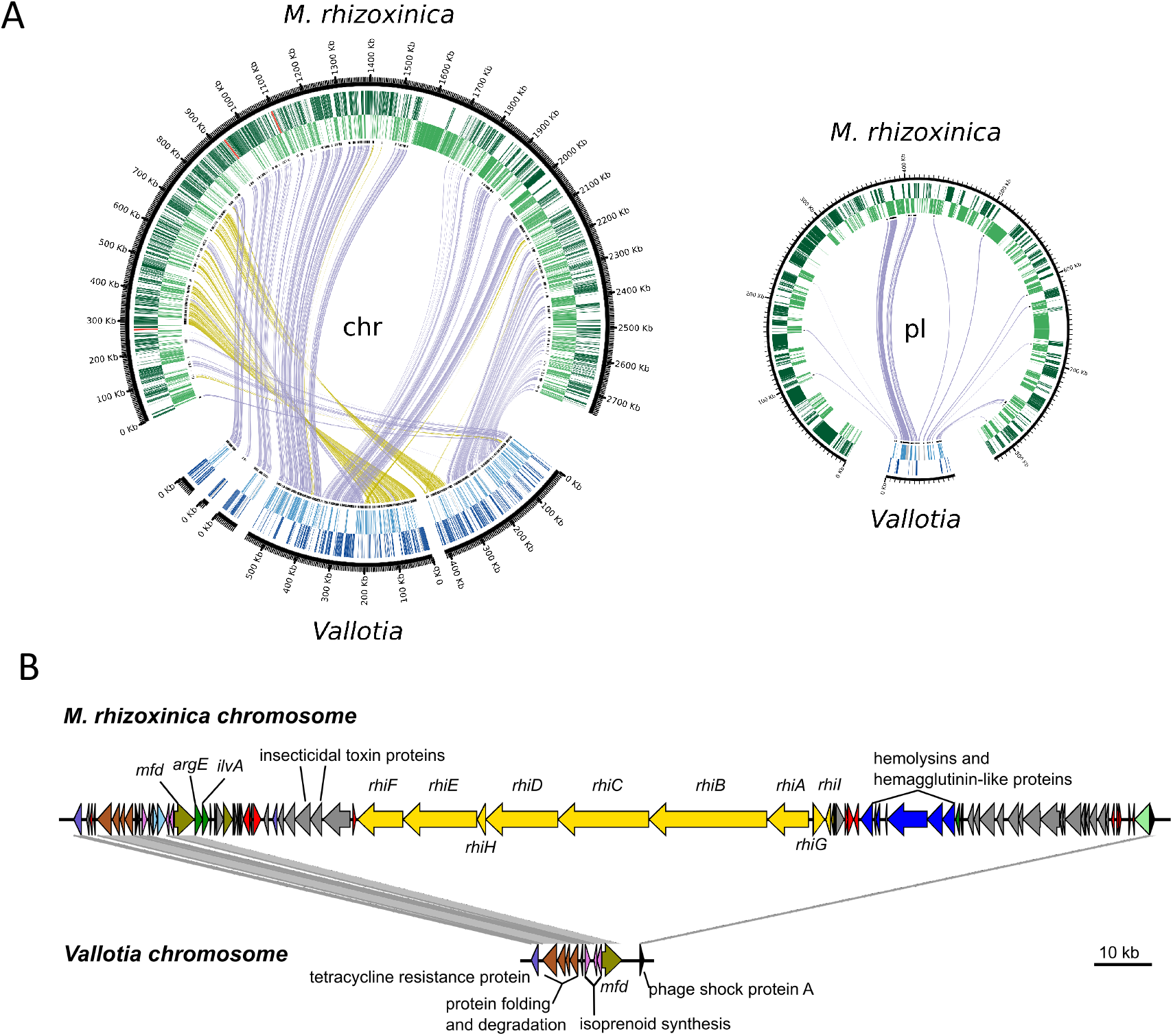
(A) Synteny between the chromosome and plasmid of *Vallotia* and *M. rhizoxinica*, an endosymbiont of *Rhizopus* fungi. The outermost and the middle rings show genes in forward and reverse strand orientation, respectively. These include rRNA genes in red and tRNA genes in dark orange. The innermost ring indicates single-copy genes shared by *M. rhizoxinica* and *Vallotia* in black. Purple and dark yellow lines connect forward and reverse matches between the genomes, respectively. The two small contigs involving the rRNA genes of *Vallotia* are not shown. **(B) Close-up of the largest deletion on the chromosome of *M. rhizoxinica* and the syntenic region on the *Vallotia* chromosome.** Genes are colored according to COG categories. Yellow: secondary metabolite biosynthesis; red: transposase; grey: unknown function; khaki: replication, recombination and repair; pink: lipid transport and metabolism; brown: protein turnover and chaperones; dark green: amino acid transport and metabolism; light green: cell envelope biogenesis; black: transcription.

Taken together, phylogenomic analyses support that *Profftia* and *Vallotia* open up novel insect symbionts lineages most closely related to free-living bacteria within the Hafniaceae and a clade of fungus endosymbionts within the Burkholderiaceae, respectively. Given the well supported phylogenetic positioning of *‘Candidatus*Vallotia tarda’ nested within a clade formed by *Mycetohabitans* species, we propose the transfer of *‘Candidatus* Vallotia tarda’ to the *Mycetohabitans* genus, as *‘Candidatus* Mycetohabitans vallotii’ (a detailed proposal for the re-classification is given in the supplementary material).

### The evolutionary link between *Vallotia* and fungus endosymbionts

#### High level of genomic synteny between Vallotia and M. rhizoxinica

The close phylogenetic relationship between *Vallotia* and *Rhizopus* symbionts offers a unique opportunity to gain insight into the early stages of genome reduction and to infer functional consequences of the partnerships of bacterial symbionts with insects and fungi, respectively. Among the *Rhizopus* endosymbionts, a closed genome is available for *M. rhizoxinica* (56). We therefore mostly focused on this fungus symbiont as a reference for comparison with *Vallotia*.

A surprisingly high level of synteny between the genomes of *Vallotia* and *M. rhizoxinica* provides further evidence for their shared ancestry. Seven contigs representing the *Vallotia* genome showed a high level of collinearity with the chromosome of *M. rhizoxinica* (Figure 3A). However, their cumulative size was only ~40% of the fungus endosymbiont chromosome. The contig that corresponded to a putative plasmid of *Vallotia* was perfectly syntenic with the larger of the two plasmids of *M. rhizoxinica* (pBRH01), although the *Vallotia* plasmid was over 90% smaller in size (72 431 bp vs. 822 304 bp) (56). Thus, the *Vallotia* plasmid showed a much higher level of reduction than the chromosome, which together with its lower G+C content and gene density suggest differential evolutionary constraints on these replicons. The observed high level of genome synteny between *Vallotia* and *M. rhizoxinica* genomes is consistent with the phylogenetic position of *Vallotia* interleaved within the clade of *Rhizopus* endosymbionts and points towards a direct evolutionary link between these symbioses and a symbiont transition between the fungus and insect hosts.

The conservation of genome structure contrasts with the elevated number of transposases and inactive derivatives making up ~6% of the fungus symbiont genome (56). Transition to a host-restricted lifestyle is usually followed by a sharp proliferation of mobile genetic elements coupled with many genomic rearrangement and gene inactivation. As seen for instance in endosymbionts of grain weevils (94), facultative and co-obligate *Serratia symbiotica* strains (95), and facultative endosymbionts, such as *Hamiltonella defensa* and *Regiella insecticola* in aphids (74). However, mobile genetic elements get subsequently purged out of the genomes of strictly vertically transmitted symbionts via a mutational bias towards deletion and because of lack of opportunity for horizontal acquisition of novel genetic elements (59,61). In contrast to the fungus symbiont, mobile elements are notably absent from the *Vallotia* genome, suggesting that they might have been lost early after the establishment of the adelgid symbiosis conserving high collinearity between the fungus and adelgid symbiont genomes.

#### Shrinkage of the insect symbiont genome

Deletion of large genomic fragments - spanning many functionally unrelated genes – represents an important driving force of genome erosion especially at early stages of symbioses when selection on many functions is weak (3,96). Besides, gene loss also occurs individually and is ongoing, albeit at a much lower rate, even in ancient symbionts with tiny genomes (62,97,98). Both small and large deletions could be seen when comparing the *Vallotia* and *M. rhizoxinica* genomes. Several small deletions as small as one gene were observed sparsely in the entire length of the *Vallotia* genome within otherwise syntenic regions. The largest genomic region missing from *Vallotia* encompassed 165 kbp on the *M. rhizoxinica* chromosome (Figure 3B). The corresponding intergenic spacer was only 3,843 bp long on the *Vallotia* genome between a phage shock protein and the Mfd transcription-repair-coupling factor, present both in *Vallotia* and *M. rhizoxinica*. Interestingly, this large genomic fragment included the large rhizoxin biosynthesis gene cluster *(rhiIGBCDHEF)*, which is responsible for the production of rhizoxin, a potent antimitotic macrolide serving as a virulence factor for *R. microsporus*, the host of *M. rhizoxinica* (88). A homologous gene cluster is also present in *M. endofungorum* and *Pseudomonas fluorescens* and it has been suggested that the *rhi* cluster might have been horizontally acquired by *M. rhizoxinica* (56,88). Rhizoxin blocks microtubule formation in various types of eukaryotic cells (88,99), thus lack of this gene cluster in ancestral *Vallotia* was likely a prerequisite for the establishment of the adelgid symbiosis. However, this large deleted genomic region also contained several transposases and many other genes, such as *argE* and *ilvA*, coding for the final enzymes for ornithine and 2-oxobutanoate productions, which were located adjacent to each other at the beginning of this fragment. The largest deletion between the plasmids encompassed nearly 137 kbp of the megaplasmid of *M. rhizoxinica* and involved several non-ribosomal peptide synthetases (NRPS), insecticidal toxin complex (Tc) proteins, and a high number of transposases among others. *M. rhizoxinica* harbors 15 NRPS genes clusters (56) in total, all of which are absent in *Vallotia*. NPRPs are large multienzyme machineries that assemble various peptides, which might function as antibiotics, signal molecules, or virulence factors (100). Insecticidal toxin complexes are bacterial protein toxins, which exhibit powerful insecticidal activity (101). Two of such proteins are also present in the large deleted chromosomal region in close proximity to the rhizoxin biosynthesis gene cluster (Figure 3B), however, their role in *M. rhizoxinica* remains elusive.

#### The Vallotia genome encodes for a subset of functions of the fungus endosymbionts

The number of protein coding genes of *Vallotia* is less than one-third of those of *M. rhizoxinica* and *M. endofungorum*, although metabolic functions are already reduced in the fungus endosymbionts compared to free-living *Burkholderia* (56). When compared to the two genomes of the fungus endosymbionts, only 53 proteins were specific to *Vallotia* (Figure S7). All of these were hypothetical proteins, and most of them showed no significant similarity to other proteins in public databases.

However, several fall within regions syntenic to the *M. rhizoxinica* genome and even retained partial sequence similarity to intact genes present solely in the fungus endosymbiont. Thus we assume that at least some of these *Vallotia* specific hypothetical proteins might rather be remnants of degrading genes and over-annotated/non-functional open reading frames than orphan genes with a yet unknown function (102,103). Four genes were present in *Vallotia* and *M. rhizoxinica* but were missing in *M. endofungorum*. These encoded for BioA, BioD in biotin biosynthesis, NagZ in cell wall recycling, and an MFS transporter. Fifteen genes, including for instance the MreB rod-shape determining protein, glycosyltransferase and hit family proteins, genes in lipopolysaccharide, lipoate synthesis, and the oxidative pentose phosphate pathway were shared between *Vallotia* and *M. endofungorum* only. The rest of the *Vallotia* genes, coding for 91% of all of its proteins, were shared among the fungus endosymbionts and the insect endosymbiont.

Comparing the genes present in both the insect and the fungus endosymbionts to those shared only by the fungus endosymbionts (Figure S8), we can infer selective functions maintained or lost during transition to insect endosymbiosis. Translation related functions have been retained in the greatest measure in the group shared by all endosymbionts. Functions, where higher proportion of genes were specific to the fungus endosymbioses, were related to transcription, inorganic ion transport and metabolism, secondary metabolite biosynthesis, signal transduction, intracellular trafficking, secretion, vesicular transport and defense mechanisms. (Most of the proteins specific to either of the fungus endosymbionts were homologous to transposases and integrases, transcriptional regulators, or had an unknown function.)

Fungus endosymbionts encode for a high number of transcriptional regulators (~5% of all genes in *M. rhizoxinica)* (56), but *Vallotia* has retained only a handful of such genes, which is a feature similar to other insect symbionts and might contribute to the overproduction of essential amino acids (62,104).

*M. rhizoxinica* is resistant against various β-lactams and has an arsenal of efflux pumps which might provide defense against antibacterial fungal molecules: the latter might also excrete virulence factors to the fungus cytosol (type I secretion) (56). Besides, *M. rhizoxinica* has several genes for pilus formation, adhesion proteins, and type II, type III, and type IV secretion systems which likely have a central role in host infection and manipulation in the bacteria-fungus symbiosis (56,105,106). However, all of the corresponding genes are missing in *Vallotia* thus neither of these mechanisms likely plays a role in the operation of the adelgid symbiosis. We could not even detect remnants of these genes in the *Vallotia* genome, except for a type II secretion system protein as a pseudogene. Loss of these functions is consistent with a strict vertical transmission of *Vallotia* between host generations, in contrast to *M. rhizoxinca*, which can spread also horizontally among fungi and can re-infect cured *Rhizopus* strains under laboratory conditions (86,87).

Additionally, the *M. rhizoxinica* genome encodes several predicted toxin-antitoxin systems (56). Plasmid-associated toxin-antitoxin systems can act as addiction molecules, which promote the maintenance of plasmids within bacterial populations (107). However, most of these are missing from *Vallotia*, only PasT, the toxic component of a chromosomal type II toxin-antitoxin system is present. Low levels of PasT can enhance bacterial stress resistance and growth of free-living bacteria, while high concentrations can induce persister cell formation (108). However, the function of PasT in *Vallotia* remains unclear.

## Conclusions

In most plant-sap feeding insects harboring a dual symbiotic system, typically the more ancient symbiont provides most of the essential amino acids. However, due to the ongoing genomic degradation characteristic for endosymbionts even genes essential in the symbiosis can get inactivated (3,58). These events might lead to the acquisition and fixation of an additional, younger symbiont, which can complement these lost functions. For instance, among Auchenorrhyncha, the universal ancient symbiont *Sulcia* provides seven or eight essential amino acids, while the rest is supplied by different younger co-symbionts (109). As a consequence of a host-restricted lifestyle, the genome of the newly arriving symbiont will also lose many functions even among those key in the symbiosis but present in the other resident symbiont (64,67,109). Co-obligate symbionts of *A. laricis/tardus* are both evolutionary recent bacterial endosymbionts of adelgids with moderately reduced genomes. This is following their occurrence in larch (*Profftia* and *Vallotia*) and Douglas fir *(Vallotia)* associated lineages of adelgids, which likely diversified relatively recently, ~47 and ~60 million years ago from the remaining clades of adelgids, respectively (5). However, these recently gained adelgid symbionts show a high level of metabolic complementarity and low functional redundancy in essential amino acid synthesis. Given its presence in both larch and Douglas fir associated adelgids, *Vallotia* might be the relatively older symbiont, and it can synthesize nine essential amino acids with a putative contribution of insect delivered compounds. Loss of functions in chorismate and anthranilate biosynthesis might have led to the fixation of *Profftia* in the system. *Profftia* can produce phenyalanine, but has lost its capabilities for synthetizing other essential amino acids. Host-derived compounds and partition of tryptophan biosynthesis between the co-symbionts in *A. laricis/tardus* are similar to other insect symbioses suggesting convergent evolution. However, the *Vallotia* - *Profftia* system differs from the *Annandia* - *Pseudomonas* system in *A. tsugae* where functions of the symbionts in essential amino acid synthesis are more balanced and redundant. It has been suggested that repeated replacement of symbionts among adelgids might be a consequence of periods with relaxed selection on symbiont functions due to different feeding behavior of adelgids on primary and secondary host trees – that is feeding on nutrient-rich parenchyma cells on spruce versus nutrient-poor phloem sap on alternate hosts – and multiple origins of host-alternating lifestyles (10). *Annandia*, the ancient symbiont of adelgids has lost many functions in essential amino acid biosynthesis, which could support this hypothesis (25), however the *Vallotia* - *Profftia* system does not follow this pattern.

One of the most remarkable findings of our study is the evolutionary link between the betaproteobacterial insect symbiont, *Vallotia*, and endosymbionts of *Rhizopus* fungi supported by their close phylogenetic relationships and a high-level of genomic synteny. There are many possible scenarios that could explain the origin of these symbioses. A common free-living ancestor could infect ancestral adelgids and *Rhizopus* fungi independently or developed an intracellular lifestyle in either of these hosts and got subsequently transmitted between them. We assume that a fungus-insect symbiont transition is more likely than multiple origins of these associations as the proliferation of mobile genetic elements typical in early stages of host restriction would have resulted in extensive rearrangements and a substantially different genomic structure (94,110), as seen, for instance, between very closely related *Serratia symbiotica* strains in aphids (95). Alternative scenarios are also possible, but the phylogenetic position of *Vallotia* interleaved within the clade of *Rhizopus* endosymbionts and lack of functions specific to the adelgid symbiont point towards the putative origin of *Vallotia* from the fungus endosymbionts. The origin of insect symbionts from fungus endosymbionts is, according to our knowledge, unprecedented. *Rhizopus* endosymbionts are equipped with many functions for infection and overcoming host defense. Chitinase, chitosanase, and a putative chitin-binding protein have also been found among the putatively *Sec* exported proteins of *M. rhizoxinica* (56), which besides the infection of fungi could have had a role in the transmission into an insect host. In addition, *Rhizopus* endosymbionts could be maintained in pure cultures (86), thus, at least for a limited time, they might survive also outside of their hosts in the environment. Their host, *Rhizopus microsporus*, is a plant pathogen fungus with a broad environmental distribution. Thus a potential route for acquisition of the symbionts by insects could have been via plant tissues, the food source of adelgids, similar to plant-mediated symbiont transmission observed for intracellular insect symbionts (21).

Taken together, our genomic analysis of co-obligate endosymbionts of adelgids revealed a novel path for the evolution of bacteria-insect symbioses from a clade of fungus-associated ancestors.

## Acknowledgements

We would like to acknowledge Alexander Siegl and Thomas Penz for their help in collecting adelgids and Irene Lichtscheidl-Schultz (Core Facility Cell Imaging and Ultrastructure Research, Cell Imaging Lab, University of Vienna) for the close-up photos of adelgids. This study was funded by the Austrian Science Fund (FWF) project P22533-B17. Work in the lab of M. H. is supported by Austrian Science Fund project DOC 69-B. A.M.M. was supported by a Marie Skłodowska-Curie Individual Fellowship (840270, LEECHSYMBIO) of the European Union.

