## Supplemental Material for "Evolutionary recent dual obligatory symbiosis among adelgids indicates a transition between fungus and insect associated lifestyles"

Supplementary Information

**B vitamin synthesis by *Vallotia***

*Vallotia* likely produces six B vitamins. For the synthesis of riboflavin, neither *yigB* nor *ybjI* (Haase et al., 2013) were found, however, the promiscuity of phosphatases has been documented (Kuznetsova et al., 2006). Thus another phosphatase, such as VALLOT_G_01570 and VALLOT_G_02690 both belonging to the Haloacid dehalogenase-like hydrolase superfamily, like YigB, might perform this reaction. As is the case for other endosymbionts, *panD* is missing for the *de novo* synthesis of pantothenate, thus this would occur from L-valine and β-alanine in *Vallotia*. Regarding pyridoxine, *Vallotia* lacks *pdxB* and *serC*, but alternatives such as the 'serendipitous pathways', *thiG* and an unspecific transaminase might bypass these steps (Kim et al., 2019; Oberhardt et al., 2016). *BioH* and *bioF* in biotin synthesis are missing from both symbionts, nonetheless, these are also notably absent in several symbiotic systems of aphids (Manzano-Marín et al., 2018; Manzano-Marín et al., 2020). *Vallotia* might still produce biotin, if either 8-amino-7-oxononanoate (KAPA) is imported or if these steps are taken over by the host. Finally, given their lack of *nadB* and *nadC* genes, both endosymbionts could synthesize NAD+ and NADP+ from the import of nicotinate.

**'*Candidatus* Mycetohabitans vallotii'** ***sp. nov.***

Based on the well supported phylogenetic positioning of *'Candidatus* Vallotia tarda' within the clade formed by both currently recognized *Mycetohabitans* species, we propose the transfer of *'Candidatus* Vallotia tarda' (NCBI taxonomy ID 1177213) to the *Mycetohabitans* genus. To keep the naming consistent, we propose the specific name *'Candidatus* Mycetohabitans vallotii' (va.lo.tii) in honor of the researcher Vallot, who described *A. laricis* in 1836. *'Candidatus* Mycetohabitans vallotii' strains have a rod-shaped cell and co-inhabit the cytoplasm of bacteriocytes of *A. laricis/tardus* along with *'Candidatus* Profftia tarda' (Toenshoff et al., 2012). We propose the old species-specific name be used as a strain name, as '*Candidatus* Mycetohabitans vallotii' strain tarda. Given their monophyletic origin, the transfer of other species of the '*Candidatus* Vallotia' genus to the *Mycetohabitans* genus is reasonable (von Dohlen et al., 2017; Toenshoff et al., 2012), however as multilocus sequence data are not available for those endosymbionts yet, we leave their species level re-designation open for future studies.

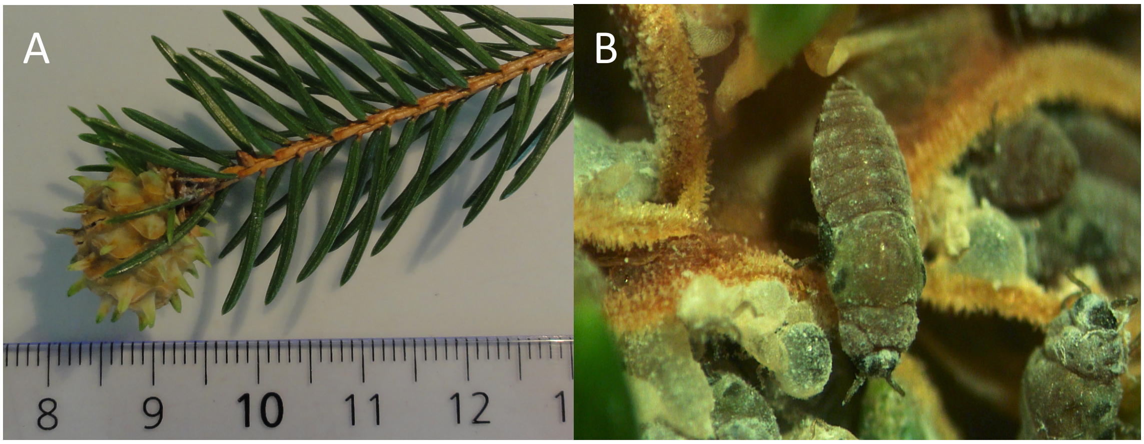


**Figure S1.** **(A)** A gall of *Adelges laricis/tardus* collected with a spruce branch. **(B)** Adelgids in an opened gall.

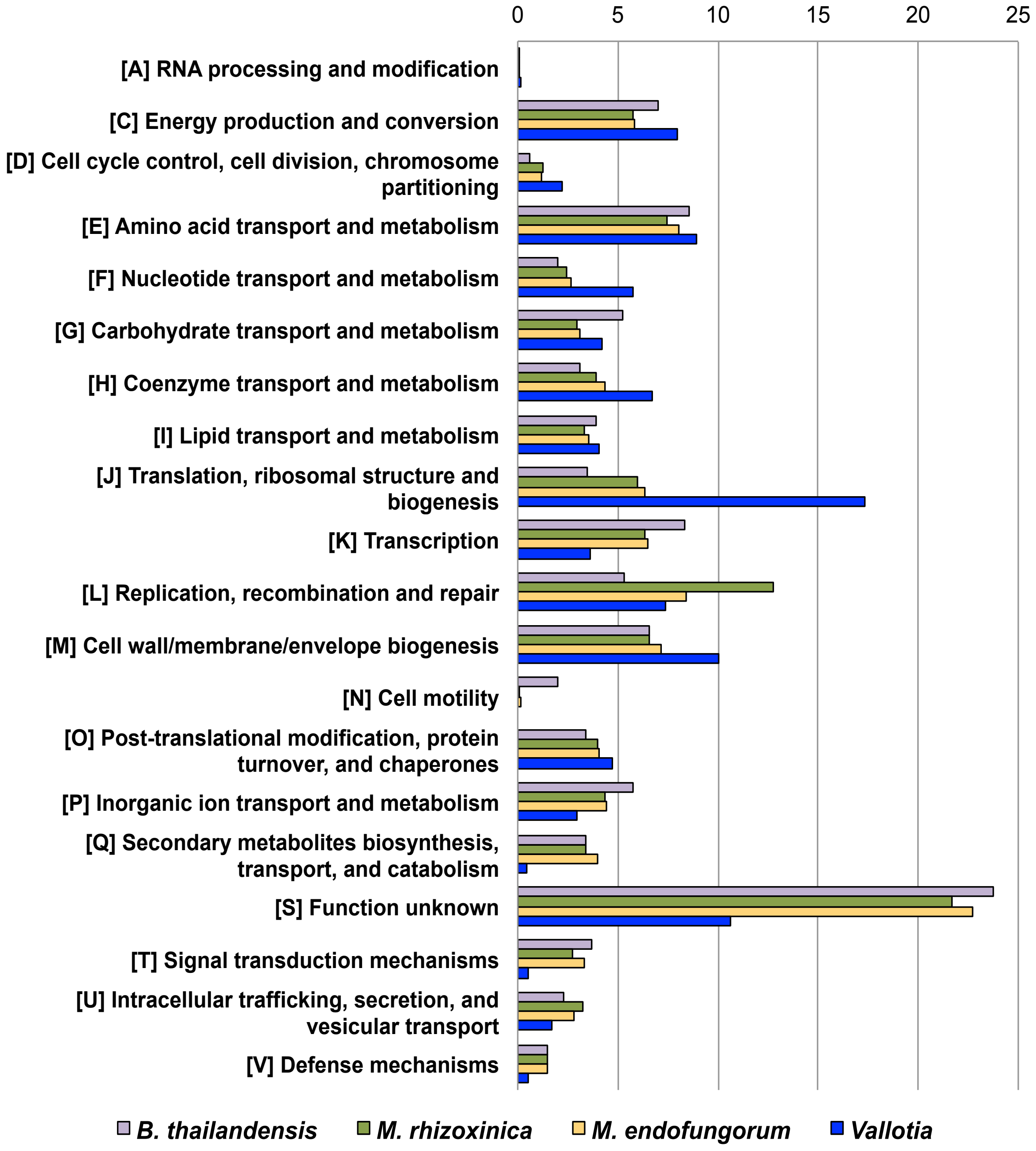


**Figure S2.** Functional reduction in *Vallotia*. Proportion (%) of genes among main functional categories according to the EggNOG classification in the genomes of *Vallotia*, related fungus endosymbionts, *Mycetohabitans rhizoxinica* [FR687359.1, FR687360.1, FR687361.1] and *M. endofungorum* [GCA_002927045.1], and a free-living bacteria, *Brukholderia thailandensis* [CP008785.1].

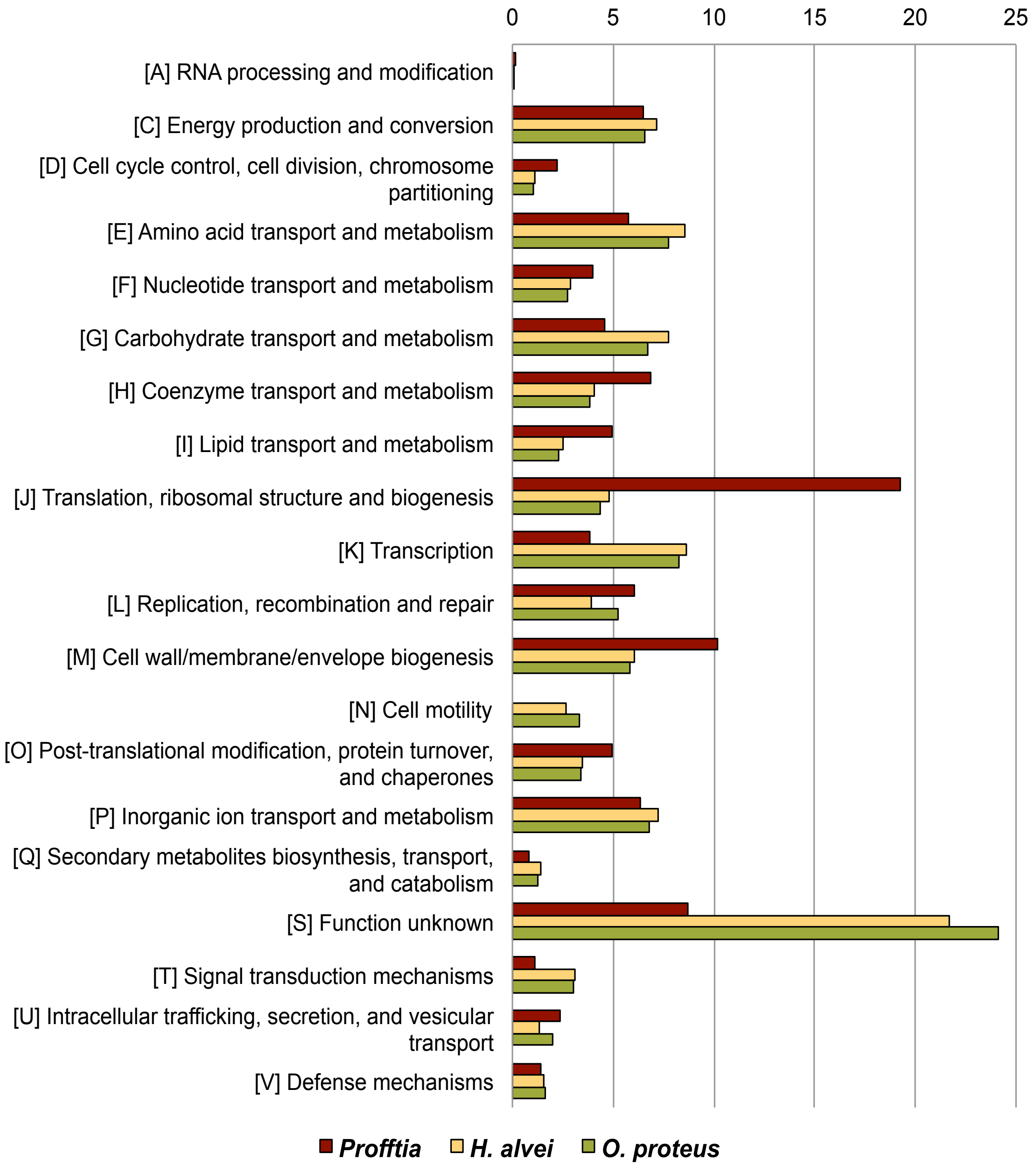


**Figure S3.** Proportion (%) of genes among main functional categories according to the EggNOG classification in the genomes of *Profftia* and related free-living bacteria, *Hafnia alvei* [CP036514.1] and *Obesumbacterium proteus* [CP014608.1].

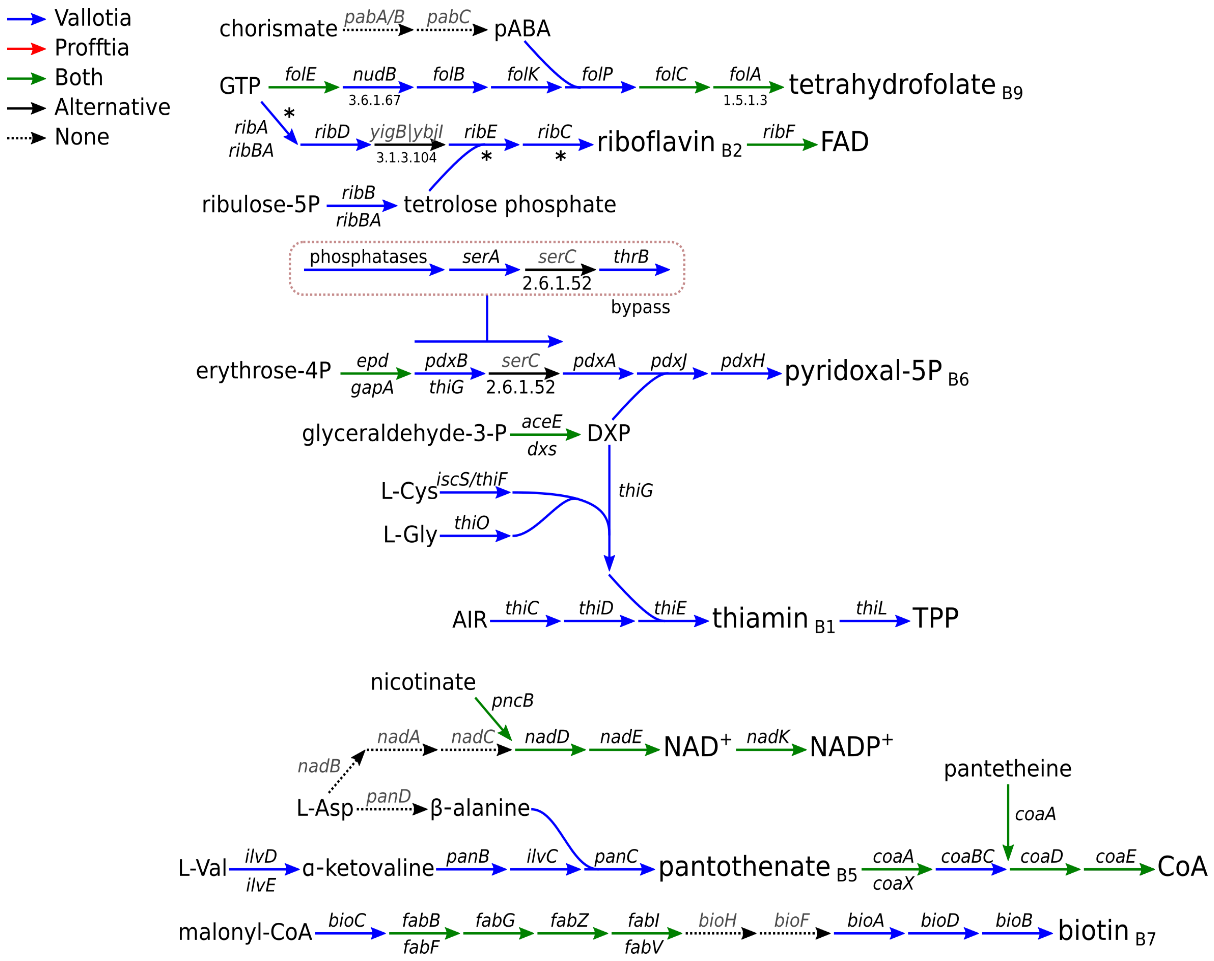


**Figure S4.** B vitamin synthesis as inferred based on the presence of genes in *Vallotia* and *Profftia.* Missing genes are shown in grey. Asterisks indicate pseudogenes of *Profftia*.

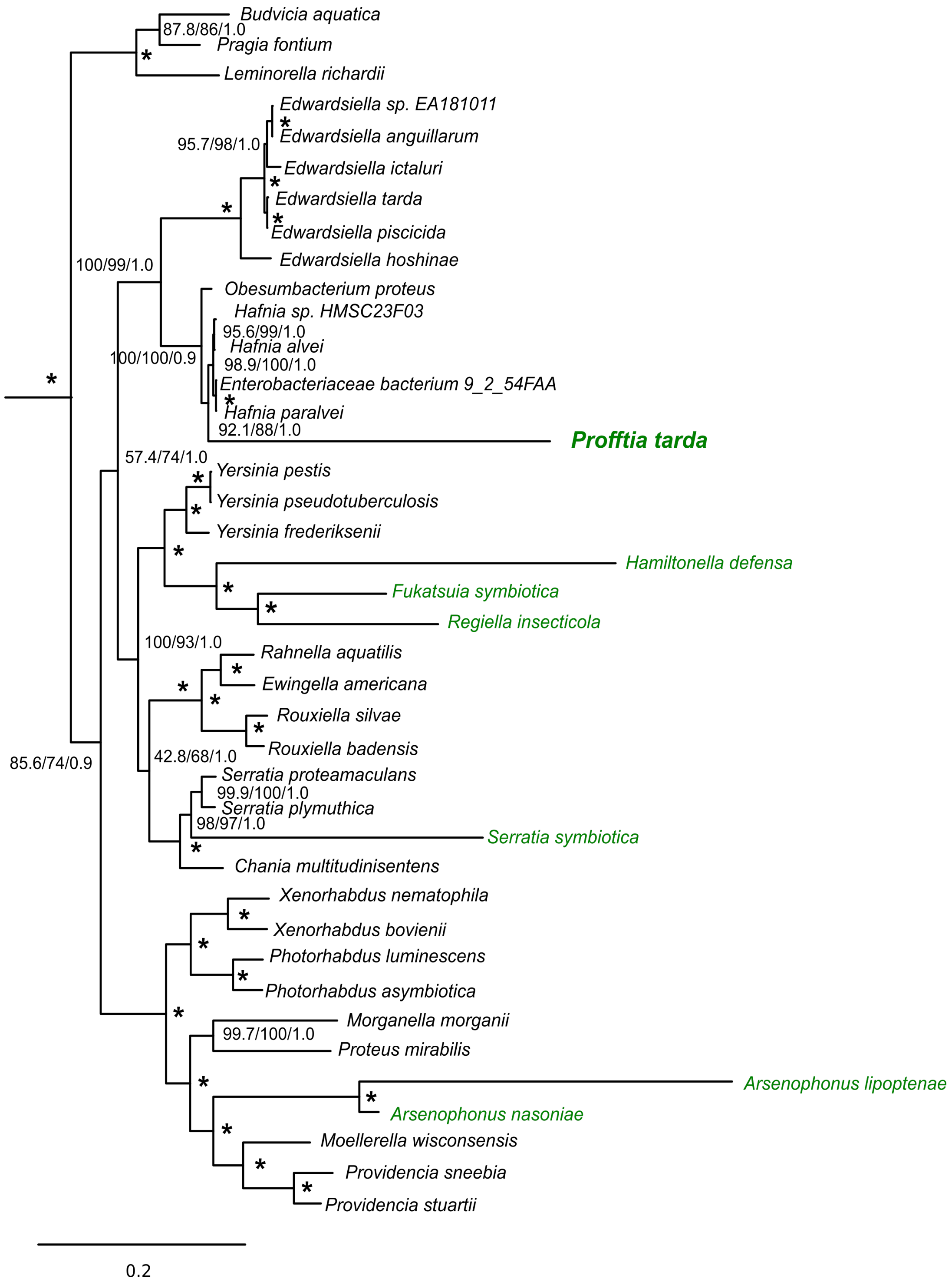


**Figure S5.** Phylogenomic analysis showing the affiliation of '*Candidatus* Profftia tarda' within the Enterobacteriales. Insect symbionts are highlighted in green. *Xanthomonas campestris* [AE008922], *Stenotrophomonas maltophilia* [AM743169] and *Pseudomonas aeruginosa* [AE004091]) were used as outgroups. Maximum likelihood (IQTREE) and Bayesian trees (MrBayes) were based on a concatenated set of 45 proteins. Maximum likelihood tree is shown. SH-aLRT support (%) and ultrafast bootstrap support (%) values based on 1000 replicates, and Bayesian posterior probabilities are indicated on the internal nodes. Asterisks stand for a maximal support in each analysis (100% / 1). The branch length leading to *Profftia* indicated accelerated evolutionary rates and was similar to those of other obligate and facultative insect symbionts included in the analysis.

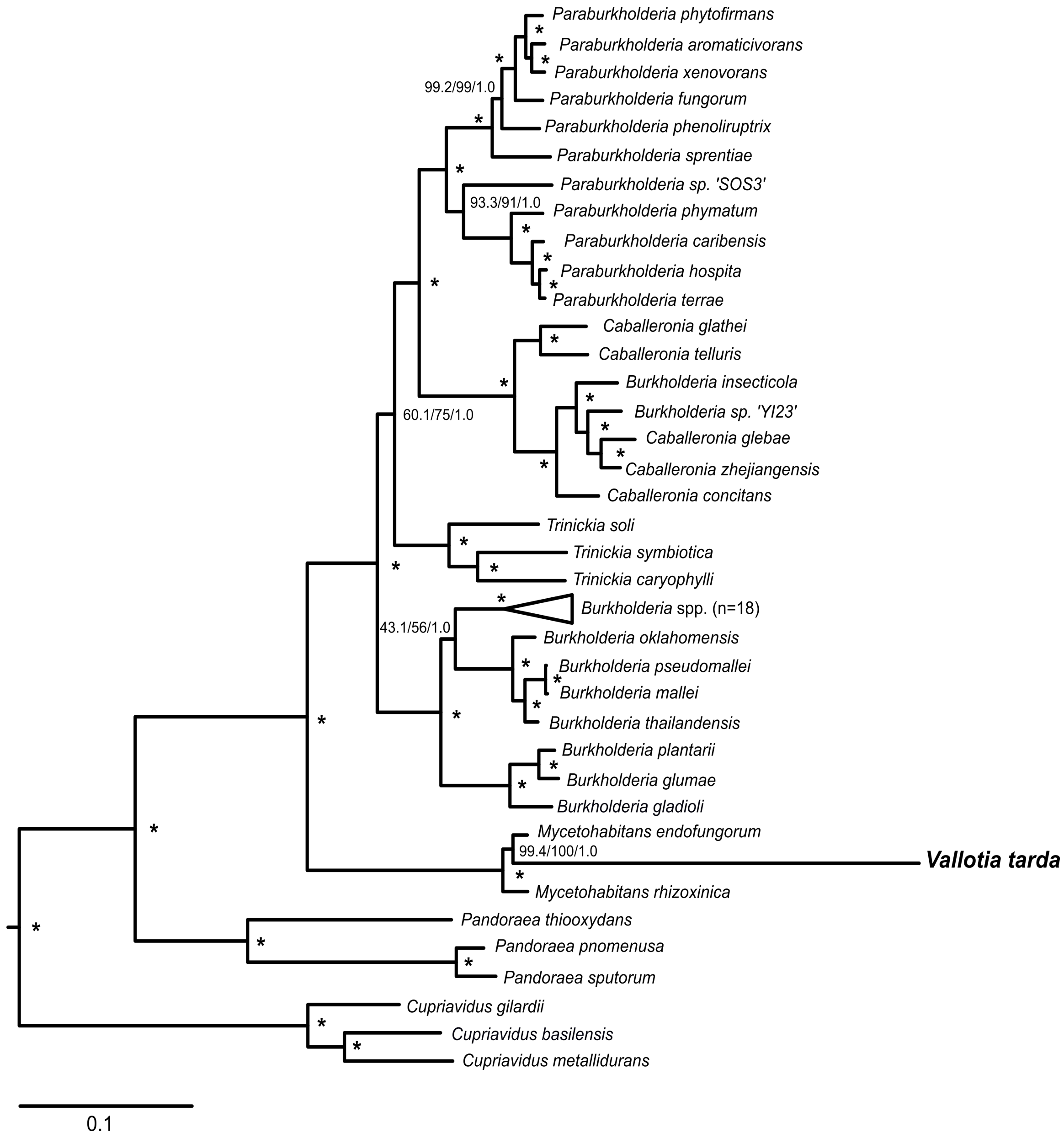


**Figure S6.** Phylogenomic analysis showing the affiliation of '*Candidatus* Vallotia tarda' within the *Burkholderiaceae*. Selected members of *Oxalobacteraceae* (*Janthinobacterium agaricidam*nosum [HG322949], *Collimonas pratensis* [CP013234] and *Herbaspirillum seropedicae* [CP011930]) were used as outgroups. Maximum likelihood (IQTREE) and Bayesian analyses (MrBayes) were performed based on a concatenated set of 108 proteins. Maximum likelihood tree is shown. SH-aLRT support (%) and ultrafast bootstrap support (%) values based on 1000 replicates, and Bayesian posterior probabilities are indicated on the internal nodes. Asterisks stand for a maximal support in each analysis (100% / 1).

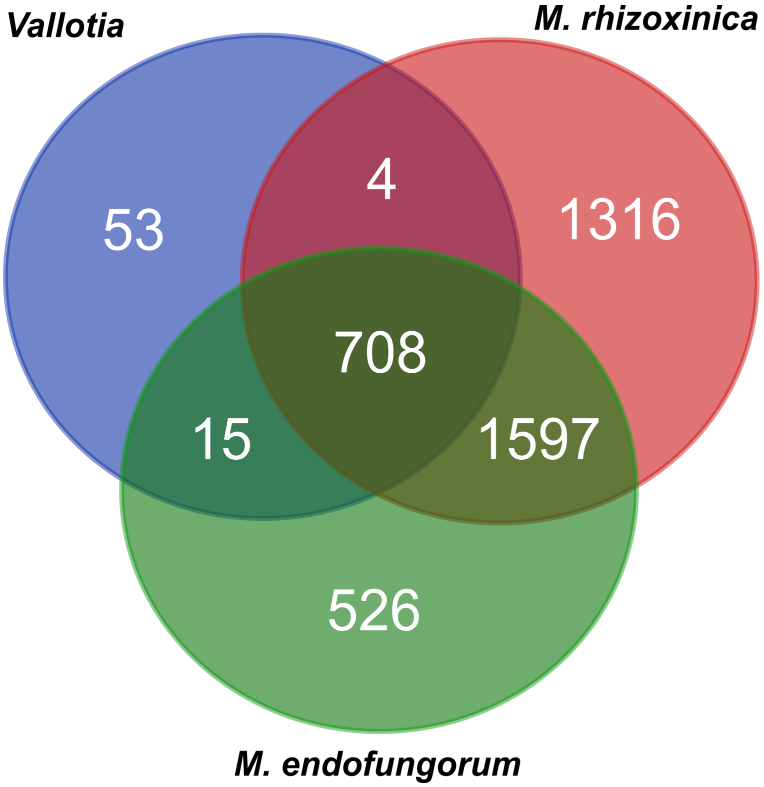


**Figure S7.** Venn diagram showing the pan-genome of the insect endosymbiont, *Vallotia*, and related fungus endosymbionts, *M. rhizoxinica* [FR687359.1, FR687360.1, FR687361.1] and *M. endofungorum* [GCA_002927045.1].

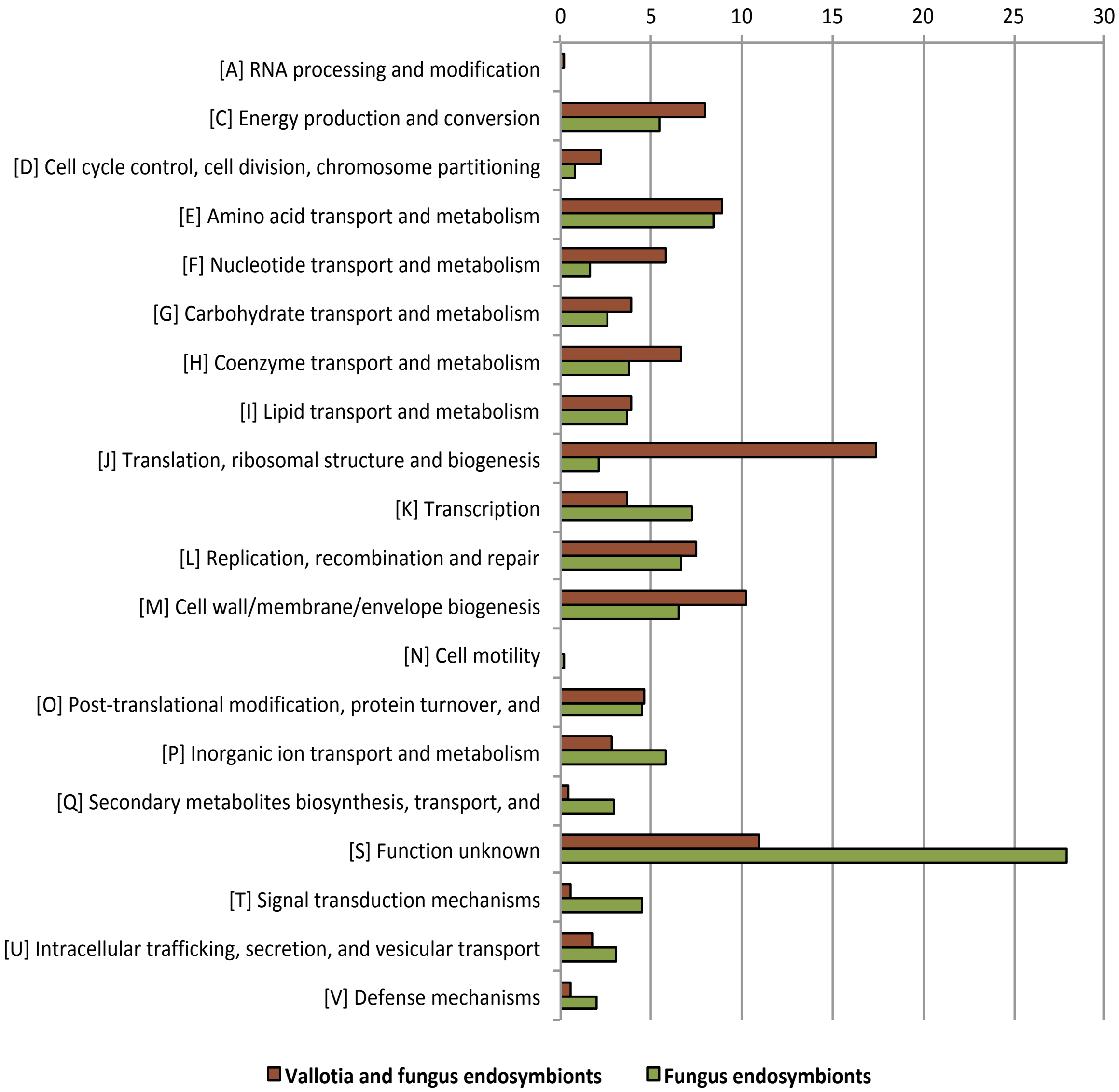


**Figure S8.** Proportion (%) of genes shared by *Vallotia* and fungus endosymbionts (shown in brown) compared to those shared only by the fungus endosymbionts (shown in green) – *M. rhizoxinica* [FR687359.1, FR687360.1, FR687361.1] and *M. endofungorum* [GCA_002927045.1] – among the main functional categories according to the EggNOG classification.
